# Assessment and Comparative Study of Biofilm Formation with frequency of Multi Drug Resistance in strains of *Staphylococcus aureus*

**DOI:** 10.1101/2021.12.16.473079

**Authors:** Kiran Fatima, Kashif Ali

## Abstract

**Background:** The study was conducted to identify the role of biofilms in the antibiotic susceptibility in the strains of *Staphylococcus aureus*. A total of 19 non repeated pus/wound swab samples from different anatomic locations and 17 samples that were previously identified as *S. aureus* and preserved in the labs were included in the study. The *Staphylococcus aureus* was identified on the basis of colony morphology, Gram’s stain, biochemical tests (catalase and coagulase tests) and molecular identification through PCR amplification.

**Methodology:** A total of 26 samples were recovered out of the 31 samples. Kirby-Bauer disk diffusion susceptibility test was used to determine the sensitivity or resistance of *S. aureus* to methicillin. Out of the 26 strains, 4 were highly resistant, 10 were moderately resistant and 12 strains were sensitive. Three different protocols (Tube Method, Congo Red Agar Method and Tissue Culture plate method) were used for the detection of biofilm formation for both resistant and sensitive strains.

**Result:** Comparative analysis of the antibiotic susceptibility and biofilm formation by different protocols showed that 70% strains that are resistant to antibiotic methicillin produced moderate-strong biofilms. 50% have produced the moderate-strong biofilms in all 3 protocols. In case of sensitive, 50% strains had produced none-weak biofilms in all 3 protocols.

**Decisions:** The strains that had zone of inhibition of close to resistance produced weak-strong biofilms but they all produced weak biofilms in CRA method. It can be concluded that the strains of *S. aureus* that have the ability to produce biofilms become methicillin resistant.

## 1. Introduction

### 1.1 Staphylococcus aureus

Staphylococcus aureus is a gram-positive, microorganism species that colonizes the anterior nares of around 20–25% human population, and 75–80% are intermittently colonized (Fenny Reffuveillle, 2017). Methicillin-resistant *staphylococci aureus* (MRSA) is one of the most important infective agents. It is the cause for several diseases from skin to serious invasive infections like respiratory disorder, infections of soft tissues, bones, heart valves, and even fatal blood disorder in humans (Gordon & Lowy, 2008).

### 1.2 *Staphylococcus aureus* Biofilms

The genetic and molecular basis of biofilm formation in staphylococci is varied. The flexibility to create a biofilm affords a minimum of 2 properties: the adherence of cells to a surface and accumulation to form multilayered cell clusters. A trade mark is that the production of the slime substance polysaccharide intracellular adhesion (PIA), a carbohydrate composed of beta - one,6 - linked N-acetyl glucosamine with part diacetylated residues, in which the cells are embedded and guarded against the host’s immune defense and antibiotic treatment. *Staphylococcus aureus* is an opportunistic infective agent that produces biofilms on medical equipment and causes respiratory disorder, meningitis, carditis, osteitis and blood poisoning. The biofilm formation by *S. aureus* involves complicated processes. The biofilm cells are controlled along and exhibit an altered composition with relevance micro-organism physiology, metabolism and gene transcription.

## 2. Materials & Methods

### 2.1 Isolation and collection of *Staphylococcus aureus*

*Staphylococci* are gram positive organisms reside in the nasal cavity and other mucous membranes as well as skin of the humans. 19 skin samples were collected by gently rubbing the tip of sterile cotton swab on the face of students at SZABIST 100 and 154 campuses. 17 samples that were preserved at campus lab were acquired.

### 2.2 Growth on Mannitol Salt Agar (MSA)

MSA is thought to be a selective media for *S. aureus. Staphylococci* can survive the high salt concentrations of MSA and thus grow without any problem. When mannitol is incited, the acid formed turns the phenol red pH indicator from **red** (base) to **yellow** (acid). All of the 19 swabs were streaked onto a small (one forth) section MSA plate using aseptic technique. The plates were left in incubator at 37°C for 24 hours.

**Figure.**
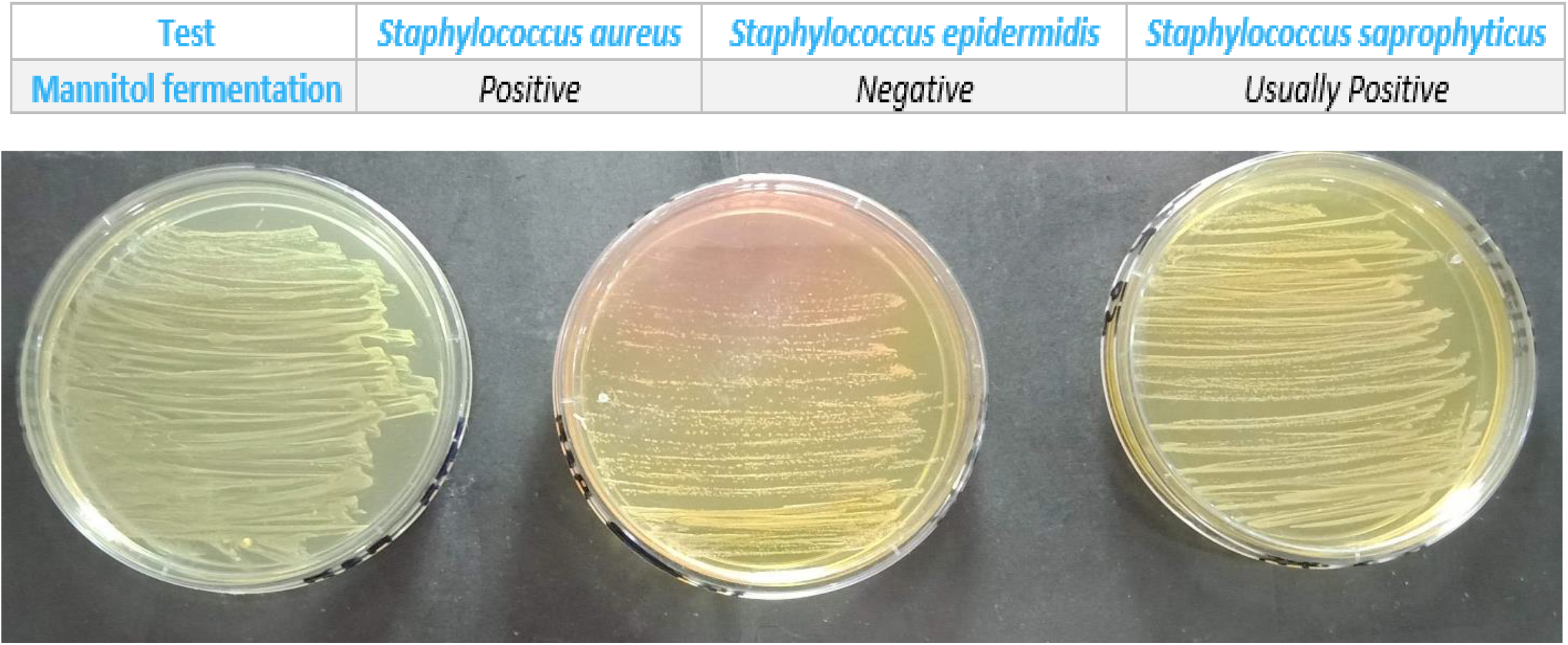
Growth of *Staph Aureus* on MSA.

### 2.3 Identification of Biochemical Characterization

#### 2.3.1 Catalase Test

*Streptococci* (that is catalase-negative) and *staphylococci* (which are catalase positive) were differentiated through catalase test. On a microscope slide small number of colonies grown on MSA were placed followed by few drops of 3% H_2_O_2_. Bubbles are produced at once if the sample contains *staphylococci*. This therefore, differentiated between *streptococci* and *staphylococci*. 31 samples were catalase positive and were separated and sub cultured. The ones that were catalase negative (did not produced the bubbles) were safely discarded.

#### 2.3.1. Catalase Test Result

**Figure.**
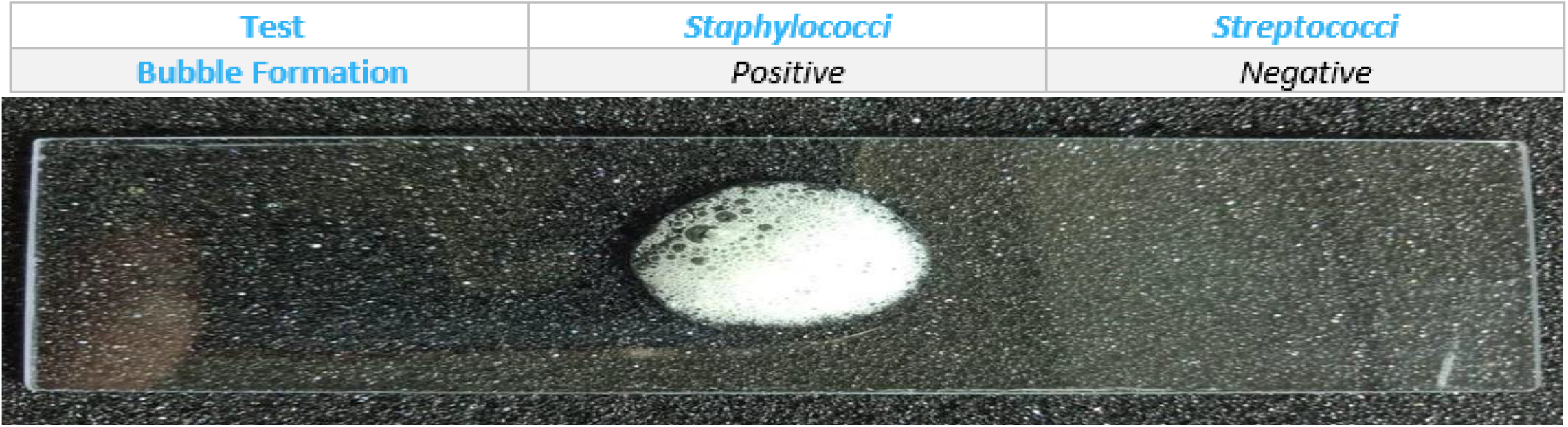
*Staphylococci* species forming bubbles when reacted with Hydrogen peroxide.

#### 2.4.1. Coagulase Test

Coagulase test was performed to differentiate the *S. aureus* from other species of *Staphylococci* as only *S. aureus* has the ability to clot the blood plasma. Hence isolating the *S. aureus* from other species of *staphylococci*. Out of the 31 samples that were tested for coagulase test 26 were coagulase positive.

#### 2.4.1. Coagulase Test Result

**Figure.**
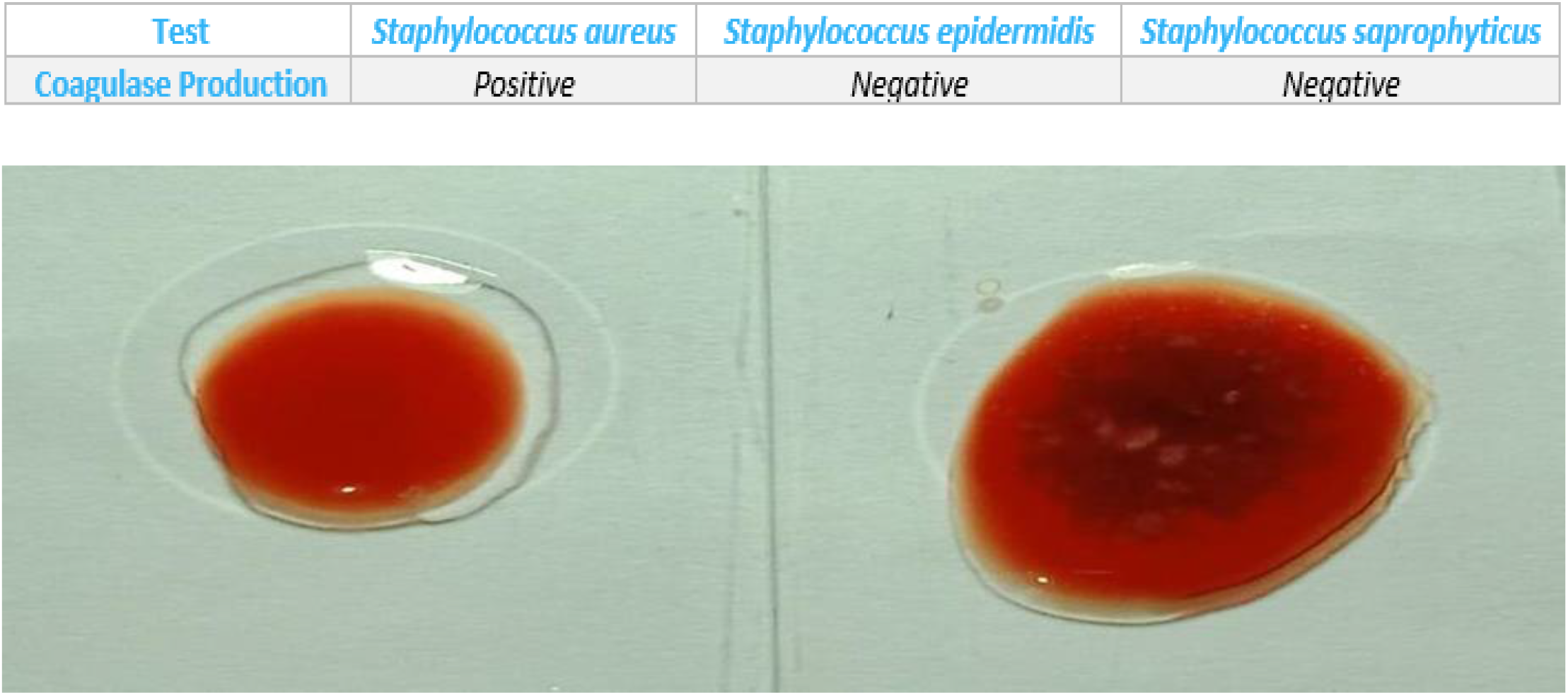
Coagulase test.

### 2.5. Testing *S. aureus* for Methicillin Resistance

Kirby-Bauer disk diffusion susceptibility test was used to determine the sensitivity or resistance of *S. aureus* to methicillin. The inhibition zone diameters were measured in millimeters using plastic (transparent) meter rule. The results were interpreted according to CLSI guidelines. An inhibition zone diameter of ≤ 13 mm was reported as methicillin resistant and ≥ 14 mm was reported as methicillin sensitive (Maj Puneet Bhatt & Kundan Tandel, 2013).

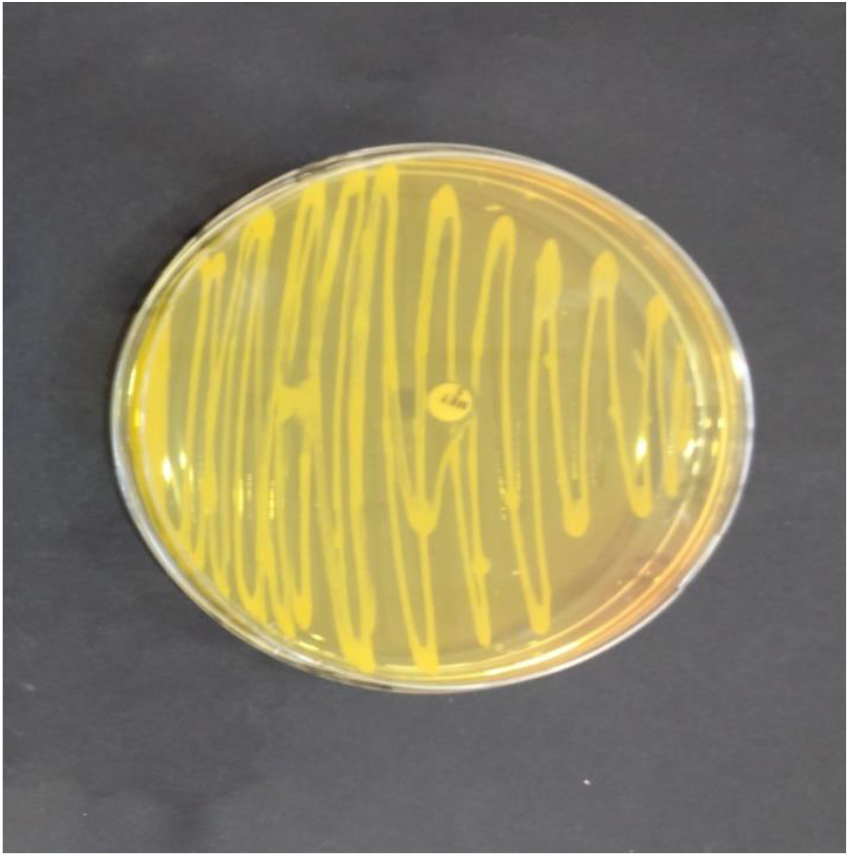

Methicillin Resistant Strain with no inhibition zone.

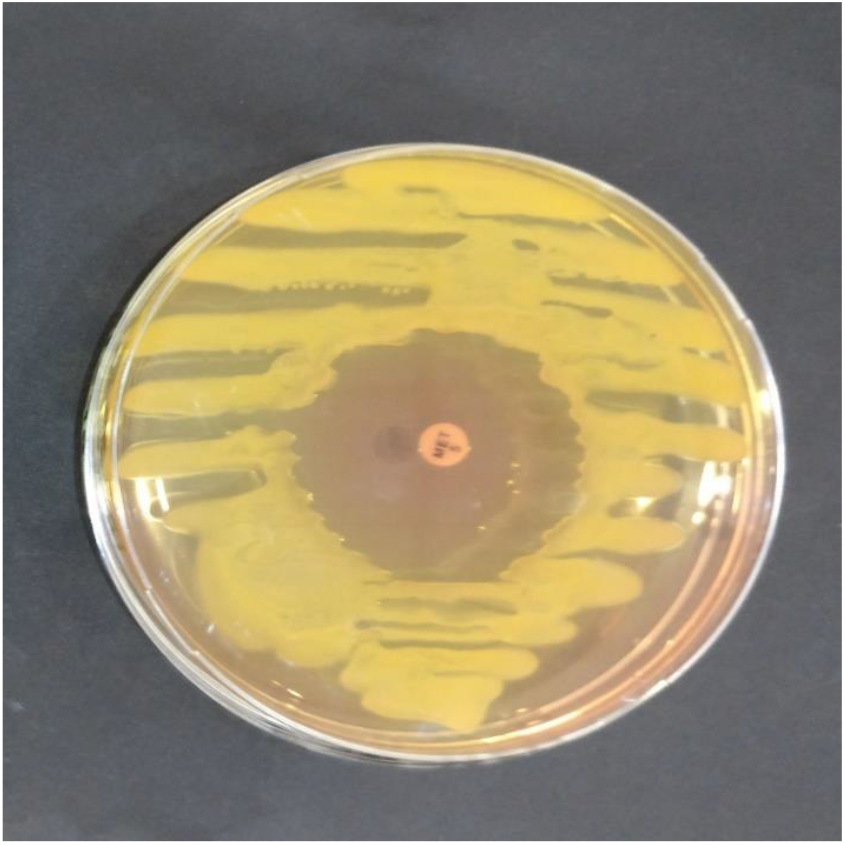

Methicillin Sensitive Strain with inhibition

### 2.6 Nature of strains tested for Methicillin resistance

| <u>Nature</u> | <u>Quantity</u> |
| --- | --- |
| Highly resistant (9 mm or less) | 4 |
| Moderately resistant (greater than 10mm ≤ 13 mm) | 10 |
| Sensitive (greater than 13 mm) | 12 |
| Total | 26 |

10 Resistant and 10 Sensitive strains were selected and were sub cultured for preservation and further use in the study **(*For the rest of the study these 20 strains were used*)**.

| <u>Strain Name</u> | <u>Zone of Inhibition diameter/mm</u> |
| --- | --- |
| **<u>Resistant</u>** |  |
| *R1* | *13* |
| *R2* | *13* |
| *R3* | *13* |
| *R4* | *13* |
| **R5** | 12 |
| **R6** | 12 |
| **R7** | 09 |
| **R8** | 08 |
| **R9** | 02 |
| **R10** | 00 |
| **<u>Sensitive</u>** |  |
| **S1** | 14 |
| **S2** | 14 |
| **S3** | 14 |
| **S4** | 14 |
| **S5** | 14 |
| **S6** | 19 |
| **S7** | 21 |
| **S8** | 22 |
| **S9** | 23 |
| **S10** | 23 |
| **Strains Used for the Study (Key; R= resistant, S= Sensitive)** |  |

### 2.7 Names Assigned and Zone of Inhibition of the Strains that were used for the rest of the Study

*To determine the relationship between the nature of the strains and biofilm production the following protocols were performed*.

### 2.9 Protocol 1: Tube method

Trypticase soy broth (TSB) was prepared with 1% glucose and poured in to the test tube. After the media was autoclaved colonies of each strains were transferred in to separate tubes and tubes placed in incubator. After 24 hours the cultures of tubes were discarded and tubes were washed with phosphate buffer saline (PBS) pH 7.3. Tubes were left to dry. 0.1% crystal violet stain was prepared and tubes were stained with it. Excess stain was discarded and tubes were washed with deionized water. The strains that produced biofilms formed a visible line on the side (wall) and bottom of the tube. **Tubes were examined and amount of biofilm formation was scored as 0-absent, 1-weak, 2-moderate, 3-strong** (Deka, 2014)

## Tube Method Results

**Figure.**
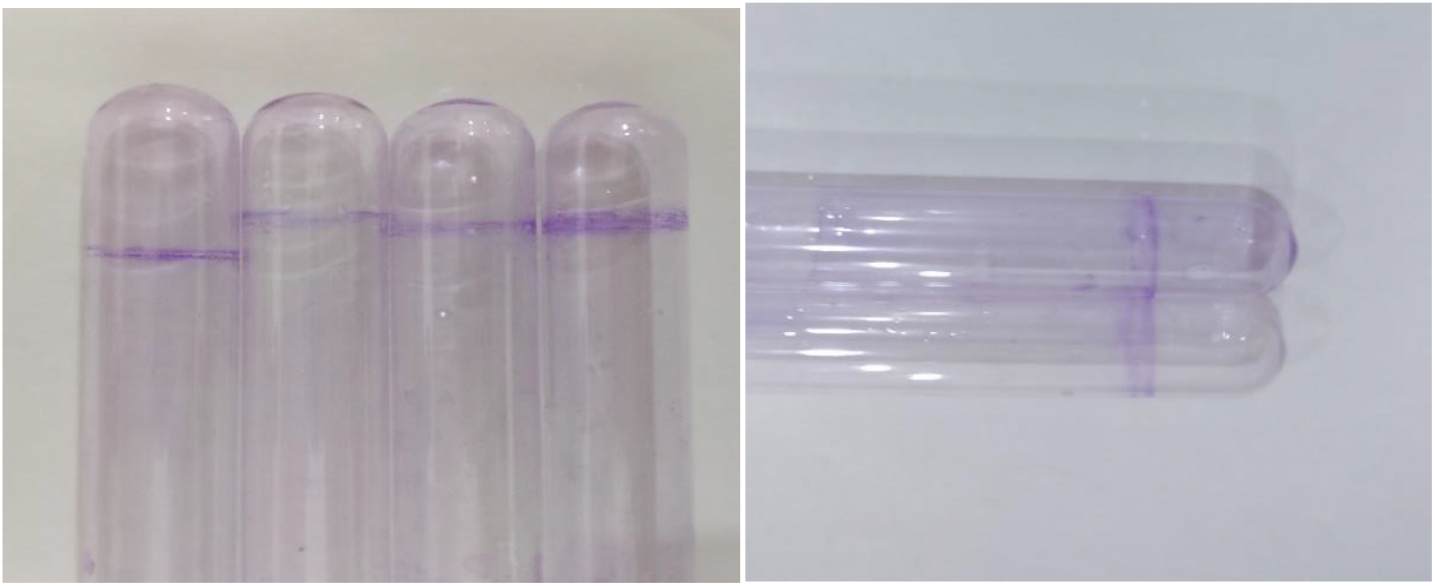
Visible film lining the wall and bottom of the tube is indicative of biofilm formation

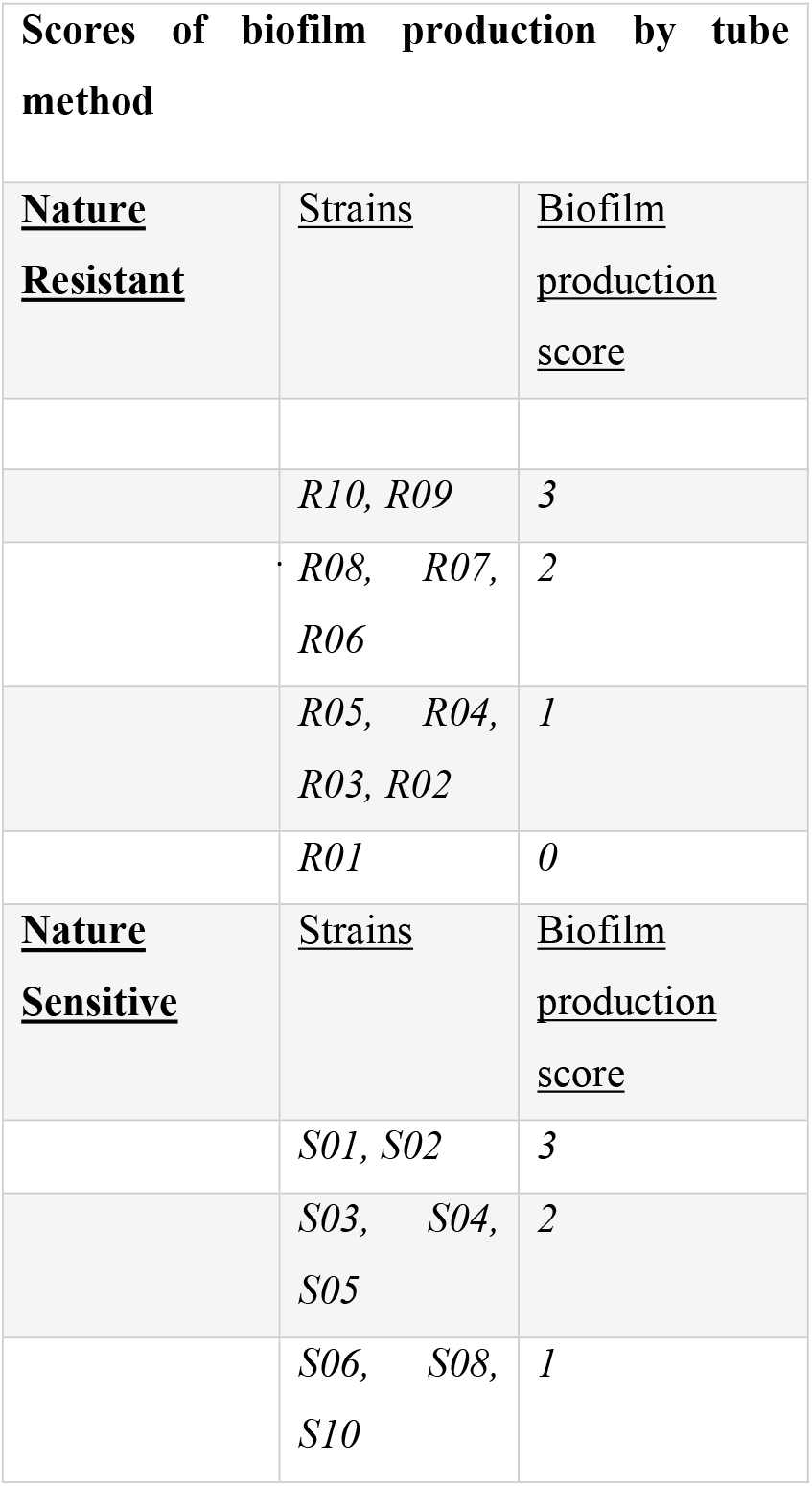
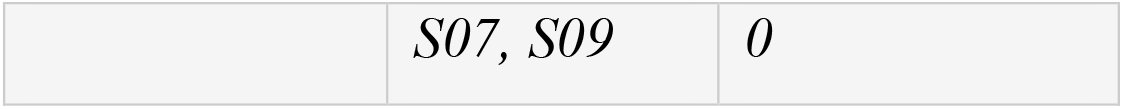

### 2.10. Protocol 2: Congo Red Agar Method (CRA)

A special medium which was mixture of Brain Heart infusion agar (37 gm/l), sucrose (5gm/l), agar no 1 (10 gm/l) and Congo red dye (0.8 gm/l) was produced. After the medium was autoclaved it was poured into the plates and strains were streaked onto the plates. Plates were incubated for 24 to 48 hours. The strains that produced strong biofilms formed black colonies and the ones that remained pink indicated weak biofilm production.

**Figure.**
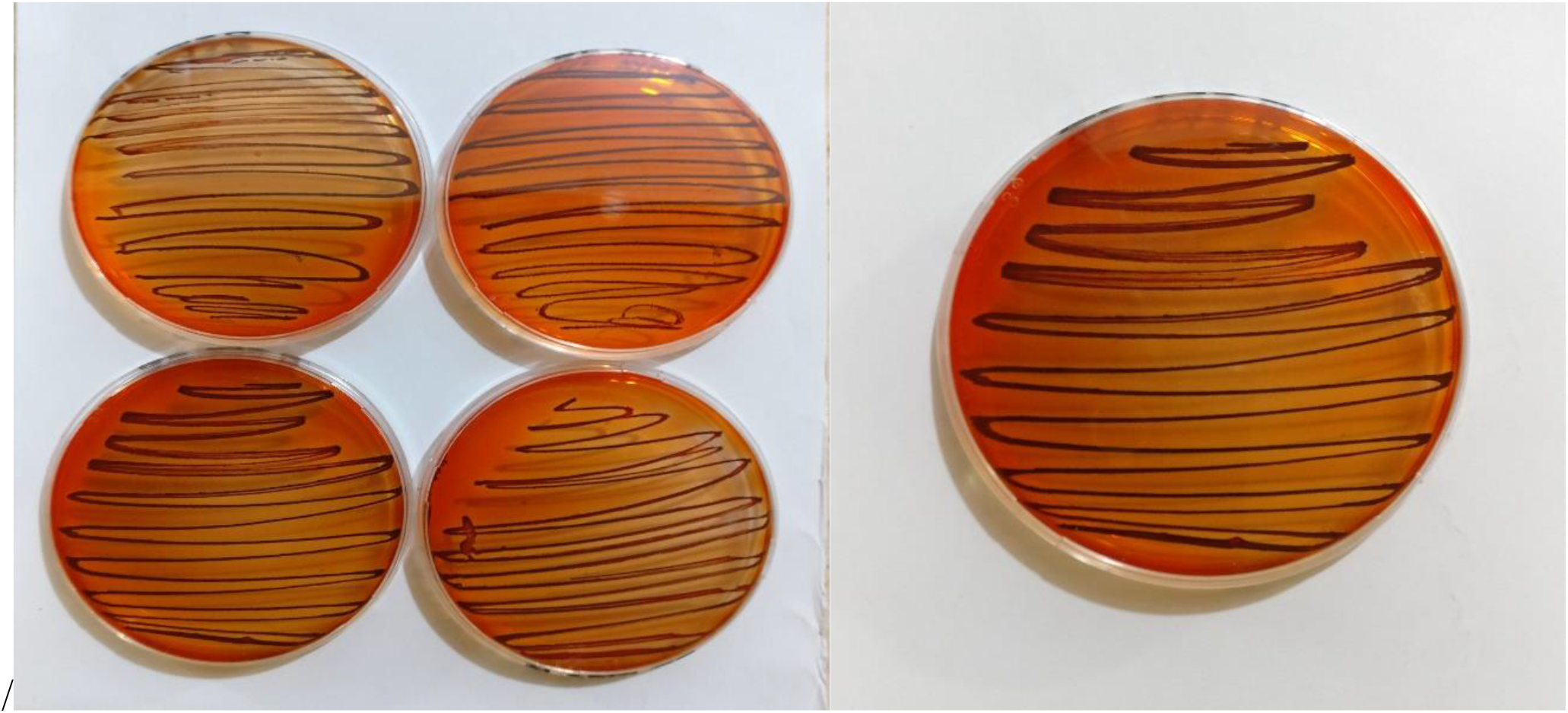
Black colonies indicating the production of strong biofilms.

**Figure.**
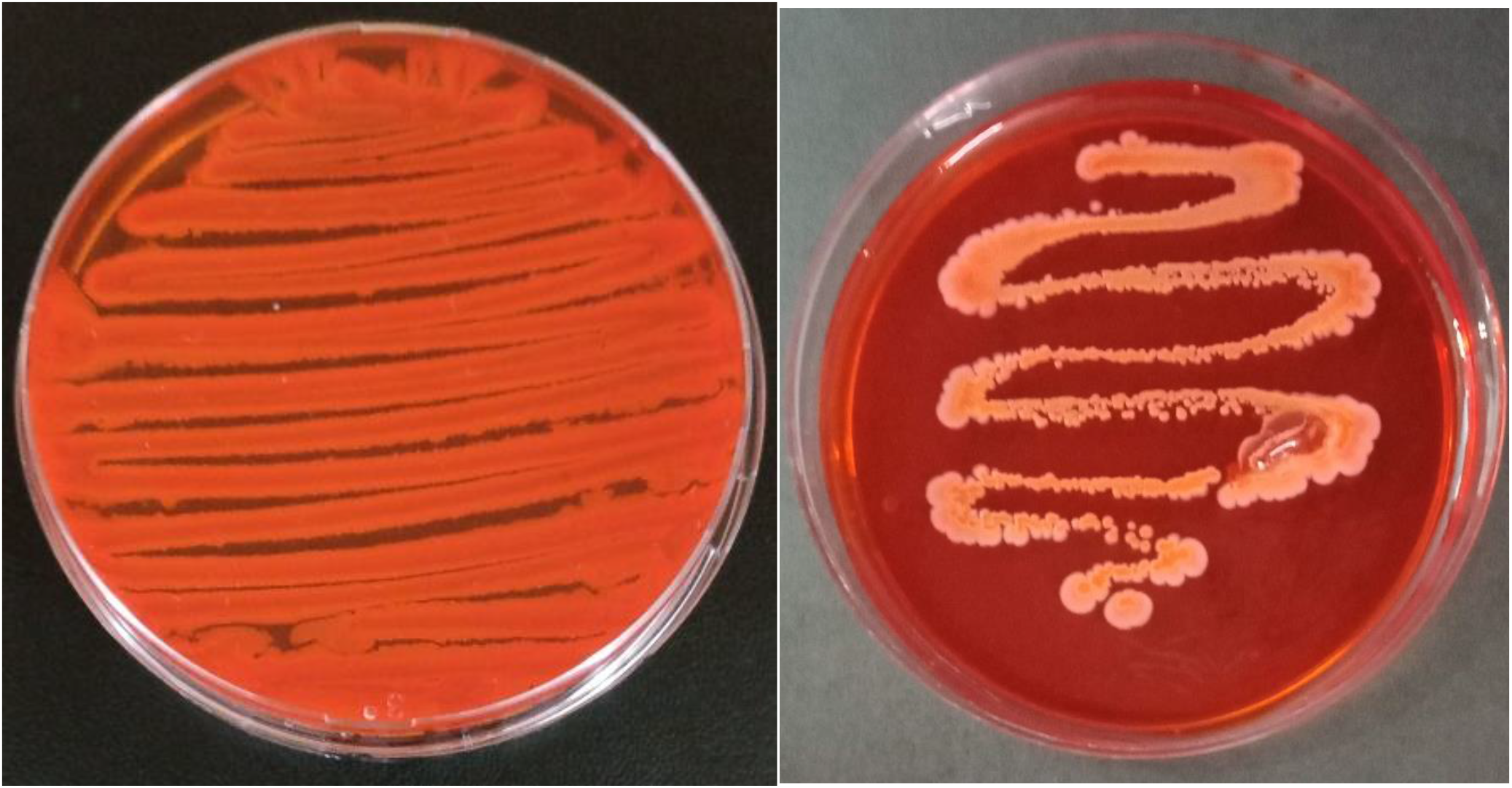
Week biofilm producing stained remained pink.

| Biofilm Production by CRA Method Results |  |  |
| --- | --- | --- |
| <u>Nature</u> | <u>Strain</u> | <u>Biofilm production</u> |
| **<u>Resistant</u>** |  |  |
|  | *R10, R09, R08, R07, R06, R04* | *Strong* |
|  | *R05, R03, R02, R01* | *Moderate-Strong* |
| **<u>Sensitive</u>** |  |  |
|  | *S01, S02, S03, S04, S05* | *Weak* |
|  | *S06, S07, S08, S09, S10* | *None* |

### 2.11. Protocol 3: Tissue Culture Plate Method (TCP)

Tryptic soy broth (TSB) was prepared with 1% glucose and poured in to the test tube. After the media was autoclaved colonies of each strains were transferred in to separate tubes and tubes placed in incubator. After 24 hours the cultures from tubes were poured into the 96 wells flat bottom micro-titre plate. Aluminum foil was used to cover the plate and plate was placed in the incubator for 24 hours. The cultures from wells were then discarded and wells were washed with PBS. 0.1% crystal violet stain was prepared and wells were stained with it. Excess stain was discarded and wells were washed with deionized water. Optical density of each well was measured at 570 nm using an automated ELISA plate reader.

**Figure.**
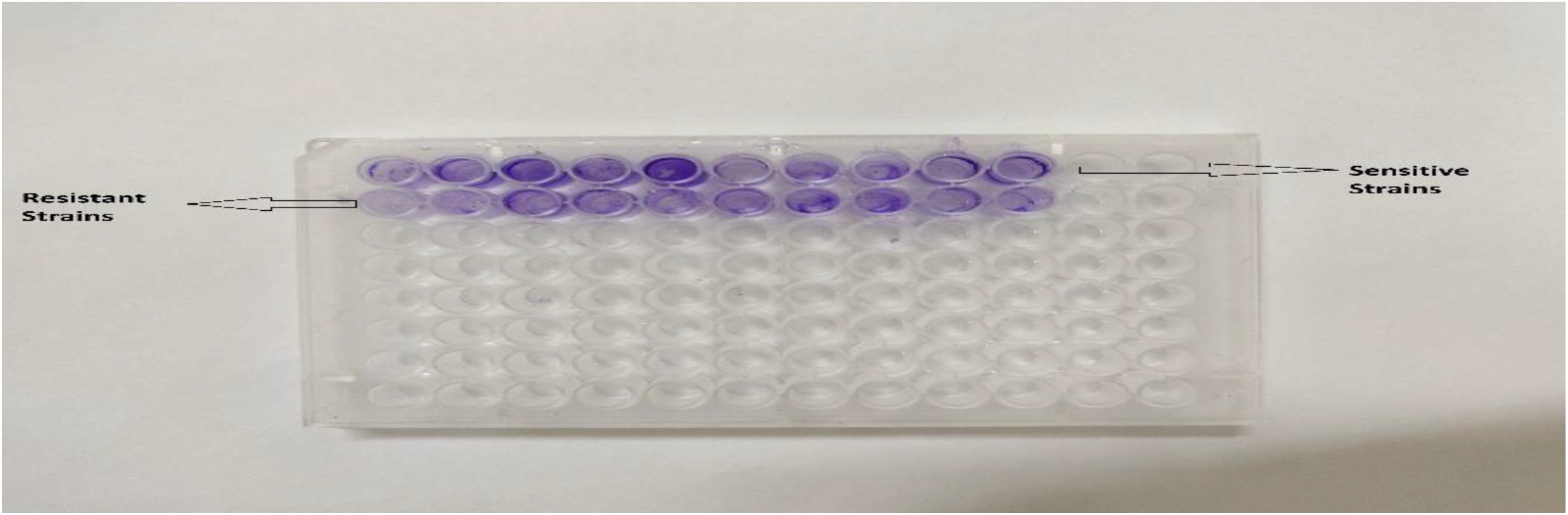
Micro-titre plate stained with crystal violet.

| Tissue Culture Plate Optical Densities |  |  |
| --- | --- | --- |
| <u>Mean OD Values</u> | <u>Adherence</u> | <u>Biofilm Formation</u> |
| $< 0.120$ | *None* | *None/weak* |
| $0.120 - 0.199$ | *Weak* | *Weak* |
| $0.200 - 0.239$ | *Moderate* | *Moderate* |
| $\geq 0.240$ | *Strong* | *High* |

Classification of Bacterial Adherence by TCP Method *(Pragyan Swagatika Panda & Dube, 2016)*

## OD Values obtained by TCP Method Results

Classification of Bacterial Adherence by TCP Method (Pragyan Swagatika Panda & Dube, 2016) OD Values obtained by TCP Method Results:
| <u>Strain</u> | <u>Mean OD Values</u> |
| --- | --- |
| <u>Resistant</u> |  |
| *R01* | *0.000* |
| *R02* | *0.155* |
| *R03* | *0.111* |
| *R04* | *0.200* |
| *R05* | *0.222* |
| *R06* | *0.227* |
| *R07* | *0.274* |
| *R08* | *0.326* |
| *R09* | *0.350* |
| *R10* | *0.354* |
| **<u>Sensitive</u>** |  |
| *S01* | *0.483* |
| *S02* | *0.304* |
| *S03* | *0.234* |
| *S04* | *0.270* |
| *S05* | *0.179* |
| *S06* | *0.130* |
| *S07* | *0.119* |
| *S08* | *0.130* |
| *S09* | *0.145* |
| *S10* | *0.172* |

## Results

Comparative Analysis of the Antibiotic Susceptibility and Biofilm formation by 3 different protocols.

| Strain | Zone of Inhibition<br>Diameter/mm | Biofilm Production By |  |  |
| --- | --- | --- | --- | --- |
|  |  | Tube Method | CRA Method | TCP Method |
| **Resistant** |  |  |  |  |
| R01 | 13 | *Absent* | *Moderate - Strong* | *None* |
| R02 | 13 | *Weak* | *Moderate - Strong* | *Weak* |
| R03 | 13 | *Weak* | *Moderate - Strong* | *None/Weak* |
| R04 | 13 | *Weak* | *Strong* | *Moderate* |
| R05 | 12 | *Weak* | *Moderate - Strong* | *Moderate* |
| R06 | 12 | *Moderate* | *Strong* | *Moderate* |
| R07 | 09 | *Moderate* | *Strong* | *Strong* |
| R08 | 08 | *Moderate* | *Strong* | *Strong* |
| R09 | 02 | *Strong* | *Strong* | *Strong* |
| R10 | 00 | *Strong* | *Strong* | *Strong* |
| **Sensitive** |  |  |  |  |
| S01 | 14 | *Strong* | *Weak* | *Strong* |
| S02 | 14 | *Strong* | *Weak* | *Strong* |
| S03 | 14 | *Moderate* | *Weak* | *Moderate* |
| S04 | 14 | *Moderate* | *Weak* | *Strong* |
| S05 | 14 | *Moderate* | *Weak* | *Weak* |
| S06 | 19 | *Weak* | *None* | *Weak* |
| S07 | 21 | *None* | *None* | *None/Weak* |
| S08 | 22 | *None* | *None* | *Weak* |
| S09 | 23 | *None* | *None* | *Weak* |
| S10 | 23 | *Weak* | *None* | *Weak* |

## Discussion

Comparative analysis of the antibiotic susceptibility and biofilm formation by 3 different protocols shows that 70% strains that are resistant to antibiotic methicillin produced moderate-strong biofilms. 50% have produced the moderate-strong biofilms in all 3 protocols which are R10 till R06. R04 and R05 have produced moderate-strong biofilms in CRA and TCP method. R09 and R10 have produced strong biofilms in all three protocols and both of them had zone of inhibitions of 2 and 0 millimeters respectively, this therefore supports the argument that the more the resistance to methicillin the stronger the biofilm is produced and perhaps more chance of producing biofilms.

In case of sensitive, 50% strains had produced none-weak biofilms in all 3 protocols. The strains that had zone of inhibition of 14 millimeters which are S01 till S05 produced weak-strong biofilms but they all produced weak biofilms in CRA method. This supports the argument that as these strains were almost near the antibiotic resistance which is less than or 13 millimeters therefore, they produced biofilms but as they were not completely resistant, they were unable to produce biofilms in all 3 protocols. Strains S06 till S10 had no or weak biofilms in all protocols and as these strains had zone of inhibition greater than 18 millimeters which is far from resistance this supports the argument that the less the resistance to methicillin the weaker the biofilm is produced or perhaps the less chance of producing the biofilms.

## Conclusion

The nature of biofilm structure and therefore the physiological attributes of biofilm organisms have inherent resistance to antimicrobial agents, no matter these antimicrobial agents are antibiotics or disinfectants. From the results obtained from the study it can be concluded that the greater the resistance to methicillin, the stronger biofilm is produced and less the resistance to methicillin the weaker the biofilm is produced or perhaps the less chance of producing the biofilms.

Furthermore, the strains of *S. aureus* that have the ability to produce biofilms become methicillin resistant. This supports the argument that biofilms play major role in providing the antibiotic resistance to bacteria.

Biofilm-producing strains of *S. aureus* pose a serious threat in health sectors. These strains of bacteria are encased in a matrix that allows them to resist and exclude antibiotics and also the host immune response. In addition to having structural barriers, the strains can rapidly undergo physiological changes such as slow growth rate and producing persistent cells. In these conditions, antibiotics fail to inhibit, kill, or eradicate these cells which are found inside the biofilm matrix. Therefore, chronic infections caused by biofilms are often difficult to treat effectively in part due to the resistance of biofilms to antimicrobial therapy. In general, antimicrobial resistance along with biofilm formation becomes an escalating and intractable problem in the health sector.

## Acknowledgement

Special thanks to Mr. Khair Bux and Ms Asma Bashir for proof reading the article, writing assistance.

## Funding

This work was supported by the Shaheed Zulfikar Ali Bhutto Institute of Science and Technology.

## Conflicts of Interest

The author declares that there are no conflicts of interest in regard to the publication of this paper.

